# Multiple Particle Tracking via Velocity Filtering (MPT-vVF): a velocity filtering framework for robust tracking moving organelles in living cells

**DOI:** 10.64898/2026.08.18.745471

**Authors:** Xinyu Liu, Zetao Fei, Ka Hei Ho, Chun Pu Wu, Jiamin Zeng, Chungwon Park, Yawen Chen, H. F. Jamie Wu, Yandong Yin, Hai Zhang, Hyokeun Park

## Abstract

Living cells are highly dynamic and densely crowded environments in which organelles such as vesicles undergo continuous motion that is essential for cellular processes. Therefore, accurate tracking of individual organelles is crucial for understanding intercellular dynamics and functions. However, precise tracking of individual organelles in living cells remains challenging due to high organelle densities, frequent particle overlap, and the coexistence of stationary and motile organelles. In particular, stationary organelles can obscure the trajectories of moving organelles, leading to tracking errors and fragmented tracks. To overcome these challenges, we developed Multiple Particle Tracking via Velocity Filtering (MPT-vVF), an unbiased, semi-automated tracking framework that incorporates a mathematically derived velocity-filtering algorithm to selectively identify and track moving organelles with high accuracy in crowded intracellular environments. MPT-vVF integrates denoising, background subtraction, and a velocity-matching detection step that discriminates true particle motion from noise based on spatiotemporal continuity, followed by robust trajectory linking. We demonstrate that MPT-vVF can accurately resolve nanometer-scale displacements of immobilized beads, highlighting its high tracking precision. We also validate the robustness of MPT-vVF by quantifying the transport of brain-derived neurotrophic factor (BDNF)-mRFP-containing vesicles in living hippocampal neurons. Furthermore, MPT-vVF reveals that exposure to 50-nm nanoplastics impairs vesicular transport, reducing both travel length and speed of BDNF-containing vesicles in living neurons. These findings establish MPT-vVF as a powerful method for quantitative analysis of intracellular organelles in crowded living cells and suggest its broad application to biophysics, cell biology, and soft matter research.

**Significance:** Accurate tracking of moving organelles in the crowded living cells remains challenging due to high particle densities, frequent overlap, and the coexistence of stationary and motile organelles. To address these challenges, we developed Multiple Particle Tracking via Velocity Filtering (MPT-vVF), an unbiased semi-automated tracking framework that uses a mathematically derived velocity-filtering algorithm to selectively identify and track motile organelles. By integrating denoising, background subtraction, and spatiotemporal velocity-matching for trajectory linking, MPT-vVF enables precise measurements of nanometer-scale displacements of immobilized beads. Moreover, MPT-vVF accurately quantifies the movement of brain-derived neurotrophic factor (BDNF)-containing vesicles and reveals transport impairment of BDNF-containing vesicles following exposure to nanoplastics. Its versatility makes MPT-vVF broadly applicable to biophysical processes, cellular transport, and particle dynamics in complex systems.

## Introduction

Living cells are highly dynamic systems that continuously sense and respond to a wide array of internal and external stimuli(1–5) to maintain homeostasis and ensure survival(6, 7). Central to this adaptability is the dynamic behaviour of intracellular organelles, which undergo constant motion, remodelling, and reorganization(8–10). These dynamic processes are essential for fundamental cellular functions(11–20) including energy production, signal transduction, and vesicular trafficking and are tightly linked to cell viability(15, 21). Consequently, disruptions in organelle dynamics are increasingly implicated in the pathogenesis of numerous diseases(15, 18, 22–30). For example, aberrant motility of organelles including mitochondria and vesicles has been reported in several diseases(15, 21–23, 26, 31, 32), underscoring the functional significance of precise spatiotemporal regulation of organelle behaviour.

Recent advances in fluorescent probes, high-sensitivity detectors, and microscopy techniques have substantially enhanced the visualization and quantitative analysis of organelle dynamics in living cells in real time(11, 12, 17, 33–36). Despite these advances, precise characterization of organelle dynamics in living cells remains challenging. The diffraction-limited resolution of conventional light microscopy, phototoxicity associated with prolonged imaging, and the crowded and heterogeneous environments hinder the accurate extraction of spatiotemporal information across diverse organelle types and length scales(12, 17, 37, 38). As a result, there is a pressing need for robust and accurate analytical methods that can quantify organelle dynamics with high accuracy from complex imaging datasets.

To address this need, numerous computational methods have been developed(10, 39–48). These methods have expanded the ability to quantify organelle dynamics, including motility, trafficking, and interactions of organelles in living cells. However, many tracking methods rely on substantial manual interventions for particle identification and trajectory correction, which are labour-intensive, susceptible to observer bias, and difficult to apply to large datasets(46, 49). Particularly, accurate tracking of organelles remains challenging in densely populated cellular environments, where frequent particle overlap, the coexistence of stationary and motile organelles, and heterogeneous motion patterns complicate particle association across consecutive frames. These factors often lead to tracking ambiguities, fragmented trajectories, and particle-switching errors, thereby limiting the accuracy and reliability of quantitative analyses of organelle dynamics.

To address these challenges, we developed Multiple Particle Tracking via Velocity Filtering (MPT-vVF), a semi-automated and robust framework for the accurate and efficient tracking of moving organelles in crowded living cells. MPT-vVF integrates denoising, background subtraction, and a velocity-matching detection step and robust trajectory linking into a unified workflow. MPT-vVF exploits motion-induced spatiotemporal patterns to suppress static background signals and to enhance the detection of moving organelles. At central innovation of MPT-vVF is the use of two mathematically grounded filters to estimate instantaneous particle velocity, providing reliable prior information during trajectory linkage, thereby minimizing mislinkage errors in densely crowded environments. We first demonstrate that MPT-vVF can accurately resolve nanometer-scale displacements of immobilized beads, highlighting its high sensitivity and localization precision. We then validate its performance in a biologically relevant system by quantifying the transport dynamics of BDNF-containing vesicles in living hippocampal neurons. Using MPT-vVF, we further show that exposure to 50-nm nanoplastics significantly impairs vesicular transport, reducing both the travel distance and speed of BDNF-containing vesicles. Together, these results establish MPT-vVF as a powerful method for quantitative analysis of organelle dynamics in crowded living cells.

## Materials and methods

### Isolation and culture of primary rat hippocampal neurons

Primary hippocampal neurons were prepared from postnatal day 0 (P0) Sprague-Dawley rat pups of either sex, sourced from the Laboratory Animal Service Centre, The Hong Kong University of Science and Technology. All experimental procedures involving animals were conducted in strict accordance with the guidelines of the Department of Health, Hong Kong Government, and were approved by the Animal Ethics Committee of The Hong Kong University of Science and Technology (AEP-2024-0041).

Hippocampal tissues were dissected from the CA3–CA1 region using Hank’s Balanced Salt Solution (HBSS; H1387, Sigma-Aldrich, USA) as previously described(11, 16, 34, 50). Tissue digestion was performed sequentially using papain (20 U/mL; LS003127, Worthington Biochemical Corp., USA) and DNase I (0.5 mg/mL; D5025, Sigma-Aldrich, USA). Following enzymatic treatment, these tissues were washed twice with HBSS and gently triturated in Neurobasal medium (A1286001, Thermo Fisher Scientific, USA) using fire-polished Pasteur pipettes until a single-cell suspension was achieved. Approximately 30000 dissociated neurons were plated onto 12-mm-diameter glass coverslips (Glaswarenfabrik Karl Hecht, Germany) pre-coated overnight with poly-D-lysine (0.1 mg/mL; P7886, Sigma-Aldrich, USA) in 24-well culture plates. Neurons were maintained in neurobasal-based conditioned medium supplemented with 2% B-27 (17504001, Thermo Fisher Scientific), 500□μM GlutaMAX-I (A1286001, Thermo Fisher Scientific), 1% penicillin–streptomycin (15140122, Thermo Fisher Scientific), and 2.5% fetal bovine serum (FBS; Invitrogen, USA). To suppress glial proliferation, 20□μM 5-fluoro-2′-deoxyuridine (FUDR; Sigma-Aldrich, USA) was added to the culture medium on days in vitro 2 (DIV2). Cultures were incubated at 37□°C in a humidified atmosphere containing 5% CO□and allowed to mature for at least 14 days. All live-cell imaging experiments were performed between DIV14 and DIV21.

### Microscopy setup and image acquisition

Imaging was performed using a custom-built widefield fluorescence microscopy system with an inverted IX73 microscope (Olympus, Japan) as previously described(16, 23). The excitation pathway included a 15× beam expander and a focusing lens to ensure uniform illumination across the field of view. Fluorescence emission from BDNF-mRFP was collected through a 100× oil-immersion objective (numerical aperture (NA) = 1.40), with immersion oil applied at the coverslip interface to maximize light collection efficiency and spatial resolution. Emitted photons were detected using an electron-multiplying charge-coupled device (EMCCD) camera (iXon Ultra 897, Andor Technology, Oxford Instruments, UK), operated via Andor Solis software. All hardware components, including the EMCCD camera, and excitation light shutter, were synchronized to enable temporally coordinated image capture.

### Beads experiments

Bead sample chambers were prepared as previously described(51, 52). Microscope slides and coverslips were sonicated sequentially in the following solutions, each for 30 minutes, followed by three rinses with Milli-Q water: first in Milli-Q water, then in acetone, next in 1□M KOH, and finally again in Milli-Q water. After sonication, the slides and coverslips were dried. The coverslips were coated with 15□µL of nitrocellulose solution and allowed to dry for 1□hour.

Preparation of fluorescent beads was performed similar to those described previously (53, 54). 100□nm fluorescent TetraSpeck microsphere beads (T7279, Invitrogen) were diluted 1:200 in PBS, and the solution was applied onto the dried coverslip. A 200□µL pipette tip was used to spread the beads uniformly. Finally, the chamber was assembled using the coated coverslip and a microscope slide separated by double-sided tape and sealed with epoxy glue. Stepping experiments using a nanometre stage were performed similar to those described previously(55, 56). The bead sample was mounted on a mechanical stage equipped with a piezo X-Y-stage (P-541.2, PI-USA) in the microscope. Regions with suitable bead density were selected. The piezo stage was programmed to move along the x-axis with a step size of 40L□nm or 60 or 80□nm every 4□seconds using PIMikroMove software. Fluorescent beads were illuminated with a 640□nm laser for signal acquisition through a 100× oil-immersion objective with NA of 1.48 (UAPON, Olympus) for total internal reflection illumination. Fluorescence signals were acquired with an EMCCD camera with an exposure time of 0.1 second via a frame transfer mode through a dichroic mirror (ZET488/640rpc, Chroma, USA) and an emission filter (ZET488/640m, Chroma) in the microscope. An additional emission filter (ET690/50m, Chroma) was inserted into the optical path upstream of the camera to selectively receive red fluorescent signals.

### Real-time imaging of BDNF-mRFP-containing vesicles in living neurons

Cultured hippocampal neurons were transfected with the plasmid construct of BDNF-mRFP by lipofectamine transfection using lipofectamine 2000 (Invitrogen) at 1000 ng/μL on DIV9 to express BDNF-mRFP in living neurons. The plasmid construct was a kind gift from Prof. Michael Silverman at Simon Fraser University. After DIV14, live-cell imaging was performed to monitor the real-time movement of BDNF-mRFP-containing vesicles. Coverslips containing transfected neurons were transferred to an imaging sample chamber. Throughout the whole imaging process, modified ACSF was continuously perfused. Fluorescence images of BDNF-mRFP were collected using 561 nm laser, a dichroic mirror (ZT561rdc, Chroma Technology) and emission filter (ET605/70m, Chroma Technology) using a frame-transfer imaging mode with an exposure time of 0.1 seconds at 10 Hz imaging.

To investigate the effects of nanoplastics exposure on the mobility of BDNF-containing vesicles, non-functionalized yellow green polystyrene nanoplastics with a dimeter of 50 nm (#17149-10, Polysciences, Inc., USA) were added to the medium in each well in a 24-well plate at a final concentration of 1 µg/mL 48-hour before imaging as previously described(50). Real-time live-cell imaging of BDNF-containing vesicles were performed at DIV14-16 using identical acquisition settings and analysed using the same procedures described above.

### Analysis

All data was analyzed by MPT-vVF. The custom-built programs written by Python and OriginPro 9.1 software (OriginLab) were used to generate histograms and fit Gaussian functions to histograms. All data are presented as the mean ± standard error of the mean (SEM). Statistical analyses were performed using OriginPro 9.1 software. Differences between groups were assessed using theMann–Whitney *U* test. A p-value of less than 0.05 was considered statically significant. Analyses were conducted under blinded conditions across multiple experimental groups.

## Results

### Multiple Particle Tracking via Velocity Filtering (MPT-vVF)

To track individual moving organelles in the densely crowded living cells robustly and efficiently, we developed Multiple Particle Tracking via Velocity Filtering (MPT-vVF). This method employs a multi-stage image processing pipeline to enhance detection fidelity and tracking accuracy of moving particles. As illustrated in Fig. 1, the workflow comprises four sequential steps: (i) denoising, (ii) background subtraction, (iii) velocity filtering, and (iv) particle linkage across frames. A key feature of MPT-vVF is that its velocity-filtering operations can be efficiently parallelized across spatial regions, temporal windows, candidate directions, and candidate speeds. GPU implementation substantially reduces computational time compared with the sequential implementation, without affecting the resulting particle detections or reconstructed trajectories.

**Figure 1.**
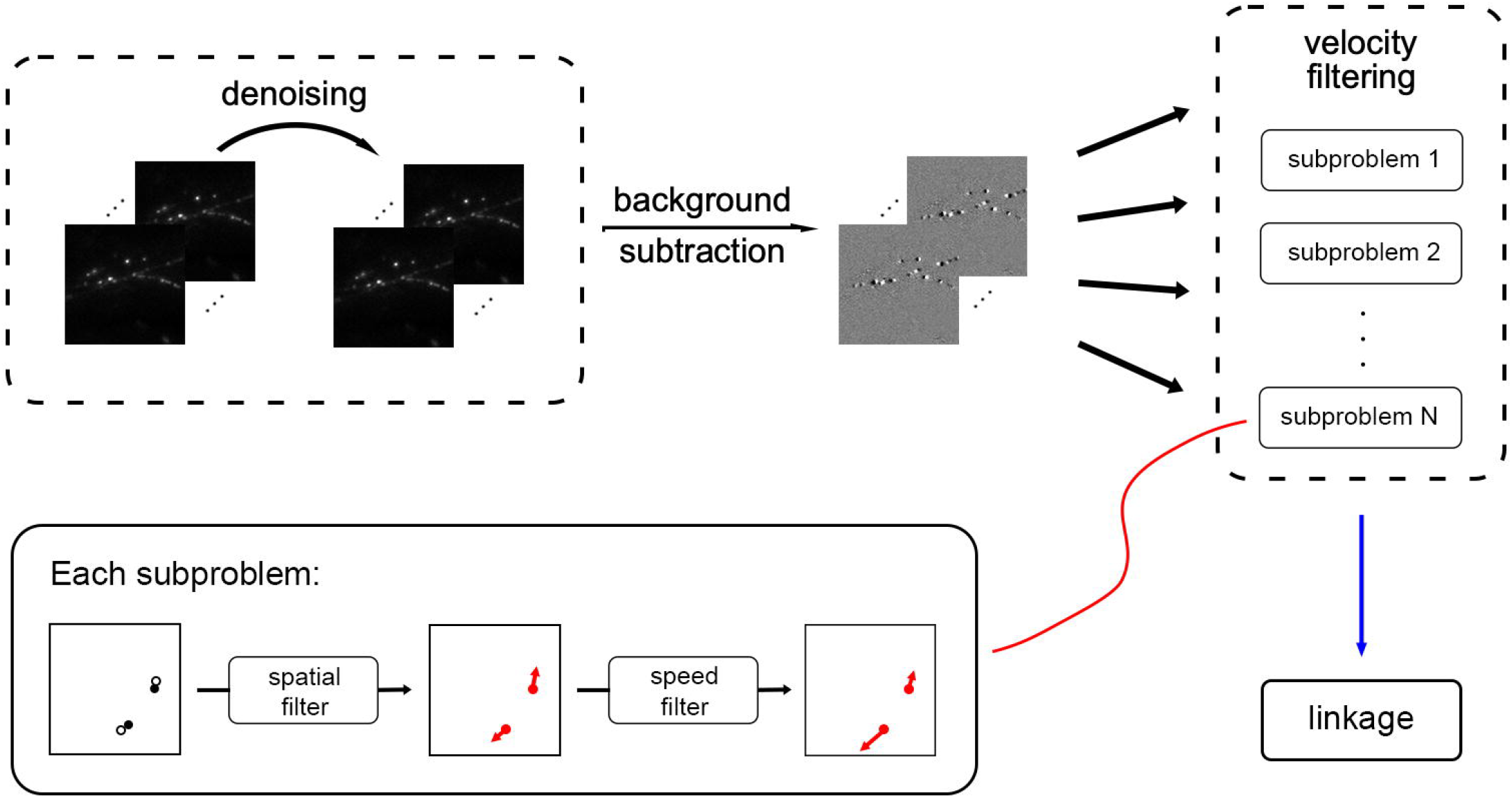
Workflow of MPT-vVF. The workflow of MPT-vVF consists of denoising step, background removal step, velocity filtering step, and linkage step. Velocity filtering process runs through parallel processing.

The denoising step as the first step is designed to suppress noise while preserving dynamics of moving organelles in the crowded living cells. Specifically, we apply a Gaussian smoothing kernel along the time dimension of the image sequence. Let *I* (*t, x, y*) denote the raw fluorescence image at time point *t*, where *t* indexes the acquisition frame number, and at spatial pixel coordinates (*x, y*). The denoised image sequence *I*_*s*_ (*t, x, y*) is obtained by convolving *I* (*t, x, y*) with a one-dimensional Gaussian kernel:

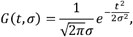

where σ is a parameter that controls the weights of the averaging. The convolution is expressed as:

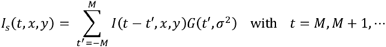

where the temporal window size is 2*M* + 1. Here, *M* is an integer chosen such that the neighboring *M* frames can be assumed to remain approximately static. In Fig. 2a, we set *M* = 3 and σ^2^ = 2.5, which effectively attenuates abrupt intensity fluctuations while retaining the temporal resolution necessary for tracking moving organelles. As demonstrated in Fig. 2b, this denoising step significantly enhances image quality by smoothing noise without compromising genuine dynamics of organelles.

**Figure 2.**
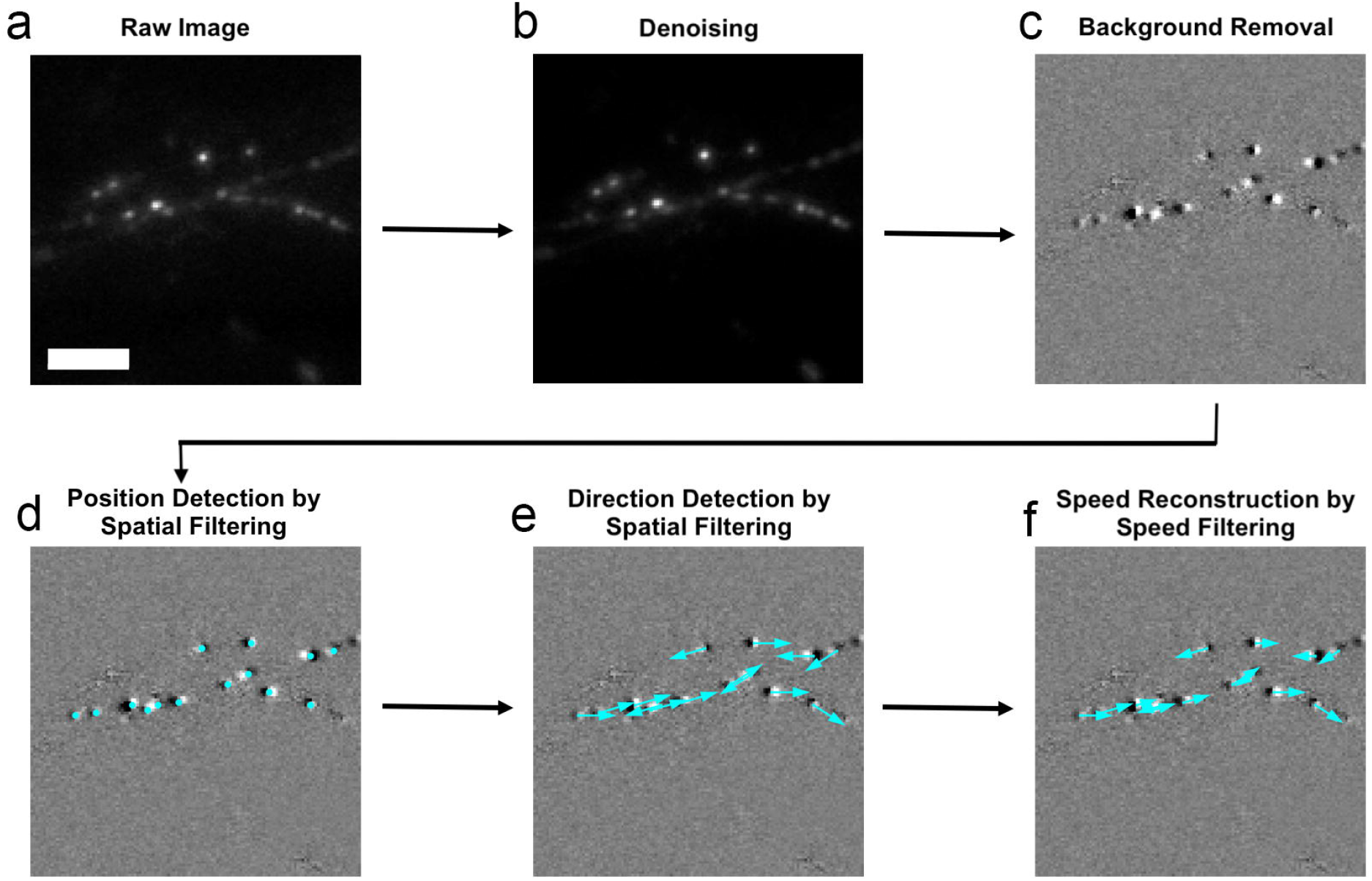
Procedures of MPT-vVF. **a**. Raw fluorescence image of BDNF-mRFP. A scale bar represents 5Lμm. **b**. Image after denoising step. A Gaussian filter was applied to the raw image. **c**. Image after background removal step. The background was subtracted from the denoised image. **d**. Image after position detection by the spatial filtering step. **e**. Image after direction detection by the spatial filtering step. **f**. Image after speed reconstruction by the speed filtering step. The integrated MPT-vVF workflow transforms raw fluorescence data into quantitative transport metrics.

To enhance the visibility of motile particles and suppress static background features from image sequence, we perform the background subtraction step by employing a frame subtraction technique. The step effectively removes signals arising from stationary particles and other nonmoving cellular structures while selectively emphasizing particle displacement between consecutive frames. As a result, moving particles are more readily distinguished from the background, facilitating subsequent velocity estimation and trajectory reconstruction. Specifically, we compute the difference between two temporally separated frames according to

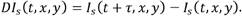

Here, *τ* denotes a frame offset chosen such that the motion of particles becomes appreciable. This step accentuates regions undergoing intensity changes, which is typically associated with particle motion, while suppressing stationary structures and persistent background signals. Consequently, the contrast of moving particles relative to the background is enhanced, improving particle detectability. By increasing the signal-to-noise ratio and reducing interference from stationary structures, this preprocessing step provides a more reliable input for subsequent velocity estimation and trajectory reconstruction. For the example shown in Fig. 2b, the time interval was set to *τ* = 5; the resulting difference image is Fig. 2c. The selection of *τ* involves a trade-off between signal strength and motion consistency. When *τ* is too small, the displacement between the two subtracted frames is limited, resulting in a weak motion-dependent signal that may be obscured by noise. Conversely, when *τ* is too large, the particle may change its speed or direction, and the positive and negative particle images may become widely separated rather than forming the gradient-like pattern assumed by the filter. Both effects can reduce the reliability of velocity estimation. Therefore, *τ* should be selected such that the expected displacement is comparable to the apparent width of the particle image.

We next apply a velocity filtering step to identify moving particles. The observation that particles in the image sequence exhibit independent motion motivates a divide-and-conquer strategy to reduce the complexity of the tracking task. Spatially, each frame is partitioned into regions with overlapping boundaries to ensure that particles moving across region borders are accurately captured without edge artifacts. Temporally, the algorithm analyzes the sequence using a sliding temporal window centered at each frame. This localized temporal evaluation allows our framework to capture instantaneous speeds of particles dynamically while preserving continuous temporal context. By decomposing the tracking task into these localized spatial and temporal subproblems, the approach enables highly efficient parallel processing. A subsequent unified linking step consistently associates the detected particles across successive framesto reconstruct continuous trajectories across the entire spatial domain.

To simplify the exposition, we initially consider the case of a single particle in the image sequence; the extension to the general case is straightforward and follows the same principle. Our proposed tracking framework employs a velocity filter consisting of two principal stages: (1) detection of the particle’s position and estimation of its velocity direction through spatial filtering, and (2) reconstruction of its speed via a dedicated speed-filtering process. Both stages are implemented using carefully designed convolutional filters. We assume the particle’s intensity profile in the image can be approximated by a Gaussian function

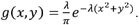

The parameter *λ* (in pixel units) specifies the spatial spread of the Gaussian function, effectively reflecting the apparent optical size of the tracked particle’s intensity profile.

After background removal, the profile of a moving particle closely resembles the gradient of this Gaussian function (illustrated by the black-and-white pair in Fig. 2c). This observation motivates the use of spatial filters derived from the directional gradient of the Gaussian. Accordingly, we introduce a set of time-independent spatial filter kernels defined as

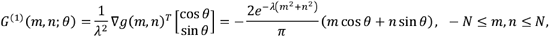

Here, 2N + 1 defines the window size of the spatial filter, corresponding to a square kernel of dimensions (2N + 1) × (2N + 1), and the factor 1/ *λ* serves as a normalization term to ensure scale consistency. *N* was selected such that the spatial filter is negligible at the kernel boundaries.

If *N* is too small, the part of the positive-negative pattern may be truncated, resulting in a weaker or distorted filter response. Conversely, if *N* is too large, additional background noise or signals from neighboring particles may be included. Because the apparent size of the pattern depends on the pixel size and optical resolution of the imaging system, *N* should be adjusted according to the imaging conditions of each dataset.

Then, we apply the spatial filter at each time point *t* as follows:

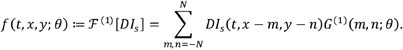

Intuitively, the spatial filter (*f*(*t, x, y; θ*)) measures the alignment between local image features around pixel (*x, y*) and the directional filter oriented at angle *θ*. A larger filter response indicates stronger evidence of a particle moving in direction *θ* at that location (*x, y*). By evaluating this function across a range of angles and positions, we simultaneously determine both the position and predominant velocity direction of the moving particle within the image sequence.

Specifically, a particle is detected at pixel location 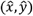 with velocity direction angle 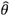 corresponding to a local maximum of *f*(*t, x, y; θ*) provided the value exceeds the predefined threshold Γ_t_. In the case of multiple particles, detections are assigned to pixel locations corresponding to distinct local maxima of the directional response over *θ*, provided that each maximum exceeds the threshold Γ_t_. Formally,

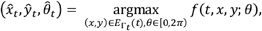

where the candidate set is defined as

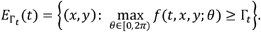

The threshold Γ_t_ is introduced to suppress spurious detections caused by fluctuations due to noise and the numerical background removal. The threshold was determined adaptivelyby the ratio of the fluorescence intensity to the noise level in the background □ subtracted image at time point *t*. Specifically, Γ_t_ was defined as

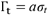

where *a* is a threshold coefficient and *σ*_t_ is the standard deviation of all pixel values in the filter-response image at time point *t*. For the image in Fig 2, *a* = 2 was used. The image-based criterion allows the threshold to adapt to variations in signal intensity and noise between frames. If Γ_t_ is set too low, background fluctuations may be incorrectly identified as moving particles, increasing the number of false-positive detections. Conversely, if Γ_t_ is set too high, dim or slowly moving particles may be excluded, decreasing the number of positive detections. The criterion of 2*σ*_t_ was selected as a practical compromised threshold to balance between suppressing noise-induced responses and retaining detectable particle signals. This detection is illustrated in Fig. 2d (location detection) and Fig. 2e (direction detection).

After determining the particle position and velocity direction, we proceed to reconstruct its speed. This step employs temporal-spatial filtering, which extends the previously introduced directional filters to incorporate temporal information. Suppose one particle has been identified at 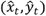 at time point *t* with velocity direction 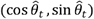. Our objective is to reconstruct the velocity (*v*_t_) of this particle at time point *t*. Let 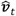 denote the estimated velocity of the particles at time *t*. Since its direction has already been estimated as 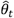, we have

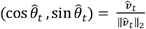

Therefore, it remains only to estimate the magnitude of velocity, or speed 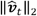. To this end, we introduce the following time-dependent speed filter kernel:

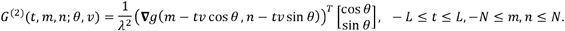

Here, *v* denotes a candidate particle speed, and *θ* is the direction of motion. The temporal window has a size of 2*L* + 1 frames. The parameter *L* is chosen such that, within the 2*L* + 1 neighboring frames over which appreciable displacement occurs, the particle trajectory can be reasonably approximated as locally unidirectional. The response for each candidate speed *v* at position (*x, y*) is computed via the speed filtering:

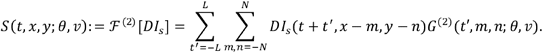

This filtering quantifies how well image intensity variations around the pixel (*x, y*) observed over a temporal window with width 2*L* + 1 align with a particle trajectory model moving at speed *v* in direction *θ*. Larger values of *S* (*t, x, y; θ, v*) indicate stronger evidence for the presence of a particle moving at speed *v* through location (*x, y*). The speed of a previously identified particle 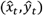 with velocity direction 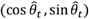 at time point *t* is estimated as the value of 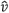 that maximizes the filtering:

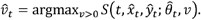

As illustrated in Fig. 2e (continuing the example from Fig. 2d), this procedure enables accurate reconstruction of particle speeds across the image sequence. In the case of multiple particles, each particle is assumed to move independently. Consequently, the position and velocity of each particle can be reconstructed individually using the same methodology. The velocity filtering step is not intended to improve spot detection. Instead, the velocity filtering step operates on the detected particles and is used to reconstruct their instantaneous velocities while enforcing temporal consistency in the tracking process. The range of velocities that can be reliably measured is bounded by both noise and moving speed. The lower velocity limit is determined primarily by the signal-to-noise ratio. Because the responses of both the spatial and velocity filters increase with particle displacement, slowly moving particles generate weak filter responses that may be indistinguishable from background fluctuations. In contrast, the upper velocity limit is constrained by the size of the local analysis region. The method assumes that a particle remains within the same local region between consecutive frames. If a particle moves too rapidly, it may exit the region before the subsequent image is acquired, resulting in inaccurate velocity estimation and reduced tracking reliability. Consequently, optimal performance is achieved when particle displacements fall within a range that is sufficiently large to generate a detectable filter response while remaining within the bounds of the local analysis window.

Following particle detection and velocity reconstruction, we link the detected particle positions across consecutive frames to obtain trajectories. Consider two consecutive image frames, *I*(*t, x, y*) and *I*(*t* + Δ*t, z, y*) Let *P*_*t*_ the set of detected particles at time point *t*, along with their associated velocities 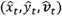. For each particle detected at position 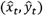 with velocity 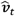, we identify its corresponding particle in the subsequent frame at time point *t* + Δ*t*. Specifically, we identify its most likely counterpart in the subsequent frame at time point *t* + Δ*t* by minimizing

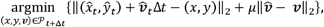

Here, the first term quantifies the Euclidean distance between the predicted particle position, which is obtained by extrapolating its current position using its estimated velocity over the time interval Δ*t*, and the actual detected positions in the next frame. The second term penalizes discrepancies in velocity between candidate matches, promoting temporal consistency in motion. The scalar parameter *µ* serves as a weighting factor that balances positional accuracy against velocity consistency. This *µ* is treated as a tunable hyperparameter and we chose *µ* = 1 in our examples. By applying this linking procedure across successive frames, we reconstruct the complete temporal trajectories of all particles.

### Measurement of nanometer-scale displacements of immobilized fluorescent beads using MPT-vVF

To assess the accuracy of MPT-vVF in resolving nanometer-scale displacements, we conducted controlled translation experiments using fluorescent TetraSpeck microsphere beads immobilized in a sample chamber mounted on a high-precision piezoelectric stage. Operating in closed-loop mode, the stage provides a positional resolution of ± 0.3 nm, enabling highly reliable sub-diffraction mechanical translations. Unidirectional displacements were applied along a single axis in predefined steps of approximately 40, 60, or 80 nm, which were selected to correspond to the physical dimensions of key subcellular structures, such as synaptic vesicles (~40 nm in diameter). Stage movements were triggered at fixed intervals (typically one step every 4 s) under software control, with the exact timing of each displacement synchronized to the image acquisition clock to ensure precise temporal alignment.

A representative fluorescence image of fluorescent beads subjected to 40 nm stage displacements is shown in Fig. 3a1. MPT-vVF detected all beads and successfully quantified their motion (Fig. 3b1). The reconstructed trajectories closely followed the imposed stepwise displacements, demonstrating high tracking fidelity. Quantitative analysis across 150 independent steps yielded a histogram with a mean step size of 40.6 ± 0.157 nm (mean ± standard error of the mean (SEM))(Fig. 3c1), in good agreement with the programmed displacement. For programmed translations of 60 nm, MPT-vVF measured a mean step size of 57.6 ± 0.310 nm (n = 147 steps; Fig. 3c2), whereas 80 nm translations yielded a mean step size of 79.6 ± 0.406 nm (n = 155 steps; Fig. 3c3). Although a slight underestimation was observed for larger displacements, the measured step sizes remained within the expected uncertainty limits of optical particle tracking at nanometer length scales. Collectively, these results demonstrate that MPT-vVF can quantify directed displacements of single fluorescent moving particles within the accuracy of tens of nanometers. The close agreement between measured and programmed step sizes across a biologically relevant range (40–80 nm), which encompasses the characteristic dimensions of synaptic vesicles, endosomes, and other organelles, validates the accuracy and reliability of MPT-vVF. These results support the suitability of MPT-vVF for quantitative tracking of organelle dynamics in live-cell imaging applications.

**Figure 3.**
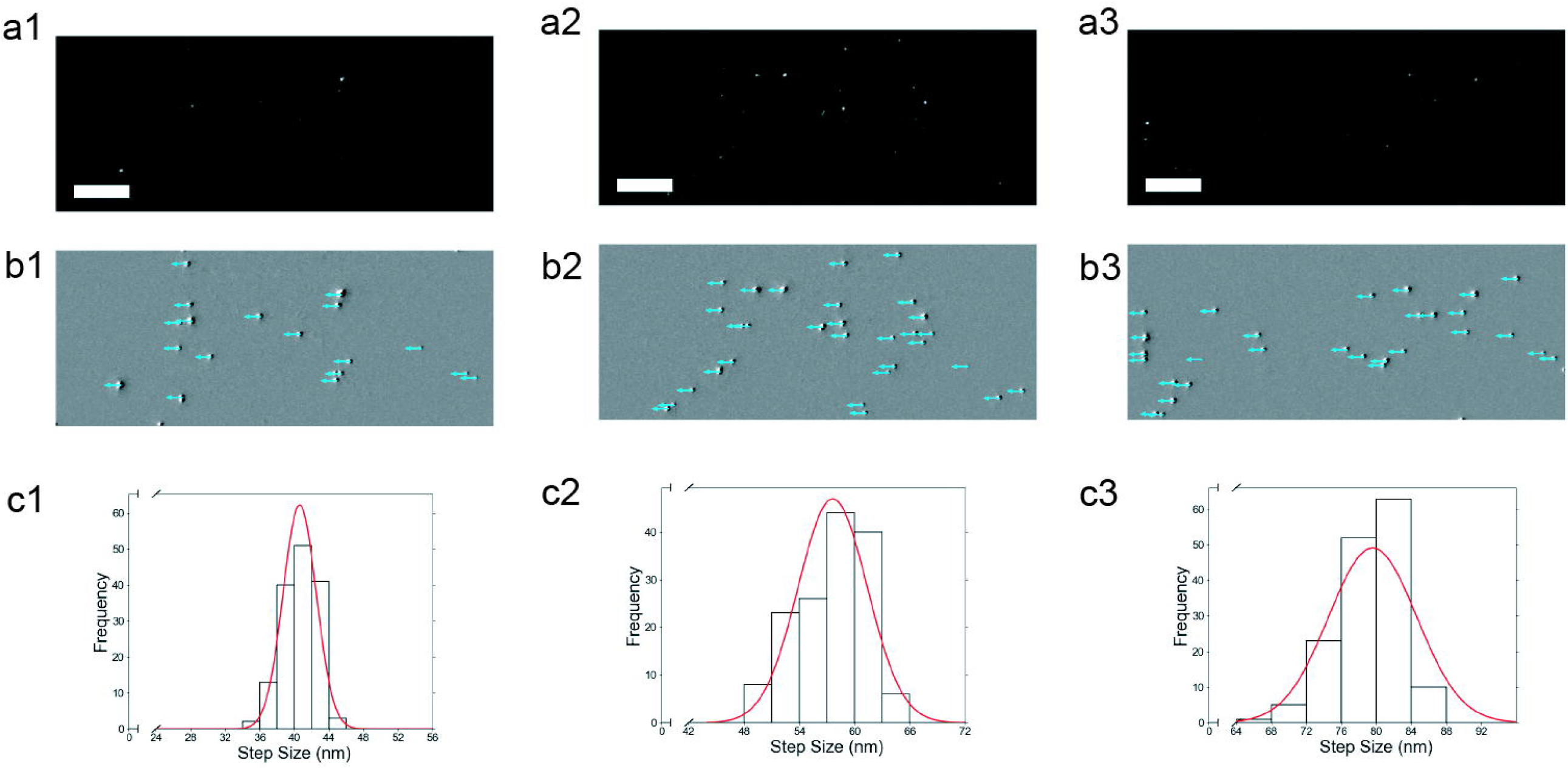
Measurement of nanometer-scale displacements of immobilized fluorescent beads using MPT-vVF. **a**. Raw fluorescence images of immobilized fluorescent beads in a coverslip on a mechanical stage moving 40 nm (a1), 60 nm (a2) and 80 nm (a3) displacements. A scale bar represents 10Lμm. **b**. Analyzed images using MPT-vVF. MPT-vVF detects all beads robustly and analyzes the motion of all beads accurately. **c**. Histogram of measured steps sizes of beads using MPT-vVF with Gaussian fitting. Measured step sizes (40.6 ± 0.157 (mean ± standard error of the mean (SEM)) nm (n = 150 steps)(c1), 57.6 ± 0.310 nm (n = 147 steps)(c2), 79.6 ± 0.406 nm (n = 155 steps)(c3)) showed good agreement with the programed steps (40 nm, 60 nm, and 80nm), demonstrating the measurement of nanometer-scale displacements of single particles using MPT-vVF.

### Tracking of single BDNF-mRFP-containing vesicles in living neurons using MPT-vVF

To assess the performance of our MPT-vVF algorithm in resolving organelle dynamics within densely crowded living cells, we transfected primary hippocampal neurons with BDNF fused to monomeric red fluorescent protein (BDNF–mRFP), a well-established reporter that selectively labels BDNF-containing vesicles(57). As shown in representative fluorescence image (Fig. 4a), BDNF–mRFP puncta were distributed throughout neurites, exhibiting marked heterogeneity in both fluorescence intensity and spatial density.

**Figure 4.**
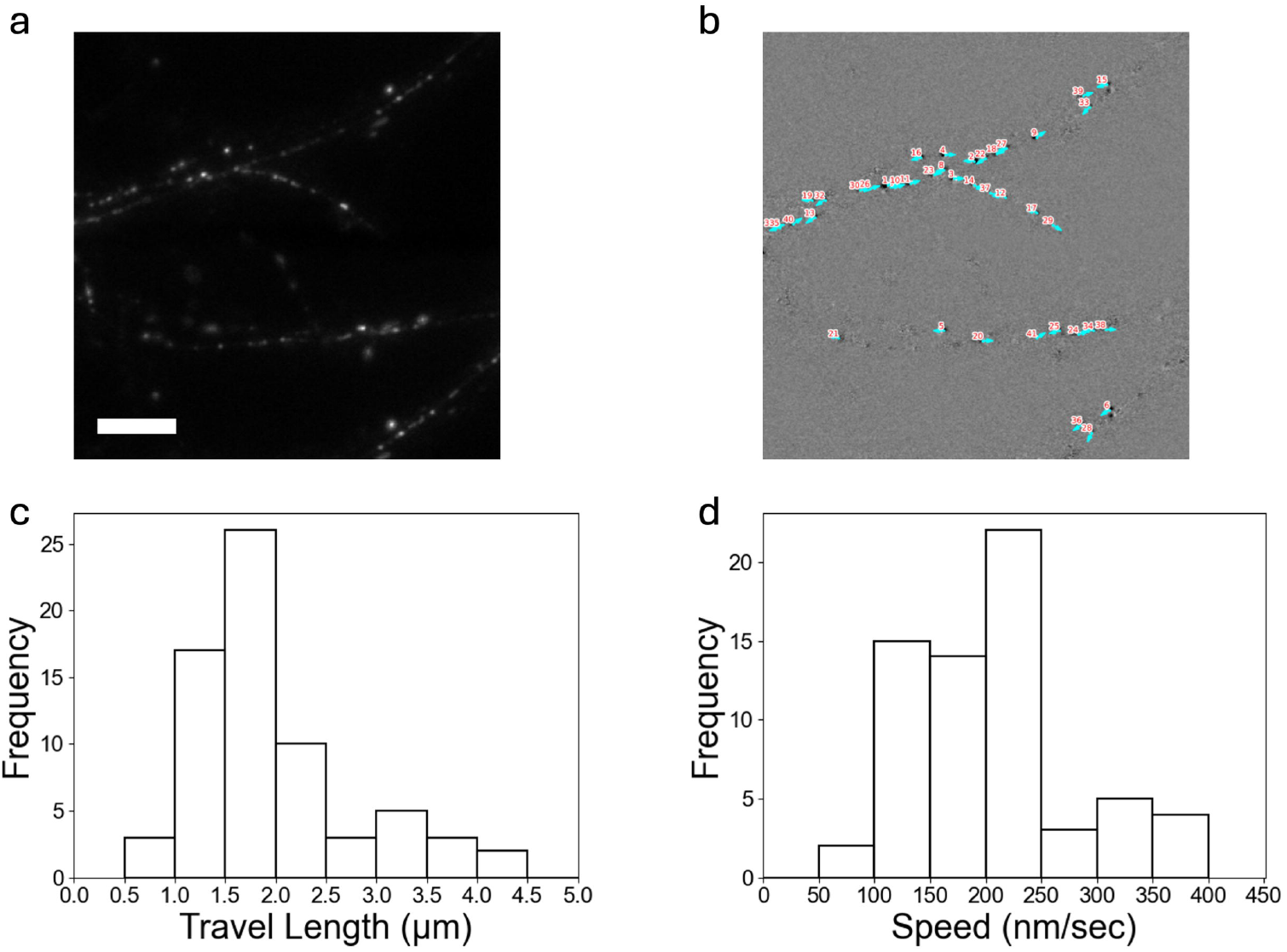
Tracking of single BDNF-mRFP-containing vesicles in living neurons using MPT-vVF. **a**. Raw fluorescence image of BDNF-mRFP-containing vesicles in living neurons. **b**. Automated vesicle detection and trajectory analysis performed by MPT-vVF. MPT-vVF accurately identified moving vesicles and analyzed the motion of all moving vesicles precisely. **c**. Histogram of travel length of single moving vesicles (n = 69 vesicles) using MPT-vVF. **d**. Histogram of speed of single moving vesicles using MPT-vVF. (n = 69 vesicles).

MPT-vVF successfully identified 41 moving BDNF-containing vesicles without manual intervention and tracked them accurately (Fig. 4b). The algorithm further enabled robust quantification of key motility parameters, such as total travel distance (= the total distance a particle covers) and speed (= total travel length/time), across extended time-lapse recordings. Quantitative analysis of 69 individual vesicle trajectories in this video of this experiment, comprising 41 moving vesicles identified in Fig. 4b and 28 moving vesicles appearing after the onset of imaging, revealed a broad distribution of travel lengths, ranging from 0.81 μm to 4.34 μm (mean: 1.98 ± 0.096 μm; Fig. 4c). Similarly, speeds varied considerably, spanning from 90 nm/s to 482 nm/s (mean: 220 ± 10.6 nm/s; Fig. 4d), similar to the previously reports of BDNF-tagged fluorescent proteins(58). This variability reflects the intrinsic heterogeneity in BDNF vesicle trafficking behaviour, likely influenced by local cytoskeletal architecture, motor protein engagement, and signalling pathways(59, 60).

### MPT-vVF reveals that exposure to nanoplastics impairs the transport of BDNF-containing vesicles in living neurons

The rapid increase in the production and consumption of plastics has resulted in the extensive accumulation of plastic waste in the environment(61, 62). Environmental plastic wastes are fragmented into microplastics and subsequently into nanoplastics(63, 64). Nanoplastics, generally defined as particles smaller than 1 μm(65, 66), pose increasing concerns due to their potential impacts on human health(65, 66). Previous studies have linked exposure to tiny plastic particles to inflammation(67, 68), reproductive toxicity(69), and cognitive deficits(70). Owing to their small size, nanoplastics cross the plasma membrane, enter cells easily and disrupt cellular function (50, 71). Recent studies showed that nanoplastics can cross the blood-brain barrier and enter the brain in mouse(72). Furthermore, nanoplastics was reported to impair synaptic functions including synaptic vesicles exocytosis and Ca^2+^ dynamics in the presynaptic terminals of living neurons(50). However, the effects of nanoplastics exposure on the transport dynamics of BDNF-containing vesicles in living neurons remain poorly understood, largely because accurate vesicle tracking is hindered by the crowded intracellular environment, where high organelle densities, frequent overlap events, and the coexistence of stationary and motile organelles complicate trajectory reconstruction. Using MPT-vVF, we reliably tracked individual BDNF-containing vesicles in primary hippocampal neurons after 48 h exposure to 1 μg/mL non-functionalized yellow-green polystyrene nanoplastics (50 nm diameter), which were characterized previously (50)). First, we measured travel length of single BDNF-containing vesicles and found that exposure to nanoplastics significantly decreased the trave length of BDNF-containing vesicles (2.17 ± 0.282 µm (*n* = 29 vesicles, *N* = 3 experiments with nanoplastics) versus 3.09 ± 0.374 µm (n = 26, *N* = 3 without nanoplastics), *p* = 0.013 (Mann-Whitney *U* test))(Fig. 5A). Moreover, exposure to nanoplastics decreased speed of BDNF-containing vesicles (241 ± 31.4 nm/s (*n* = 29, *N* = 3 with nanoplastics) versus 343 ± 41.6 nm/s (n = 26, *N* = 3 without nanoplastics), *p* = 0.013 (Mann-Whitney *U* test))(Fig. 5B). The observed decreases in travel length and speed indicate that exposure to nanoplastics disrupts the mobility of BDNF-containing vesicles in living neurons. Collectively, these results validate MPT-vVF as a powerful tool for quantitative analysis of intracellular cargo dynamics in living neurons, with utility for studying subtle alterations in vesicular transport under physiological or pathological conditions.

**Figure 5.**
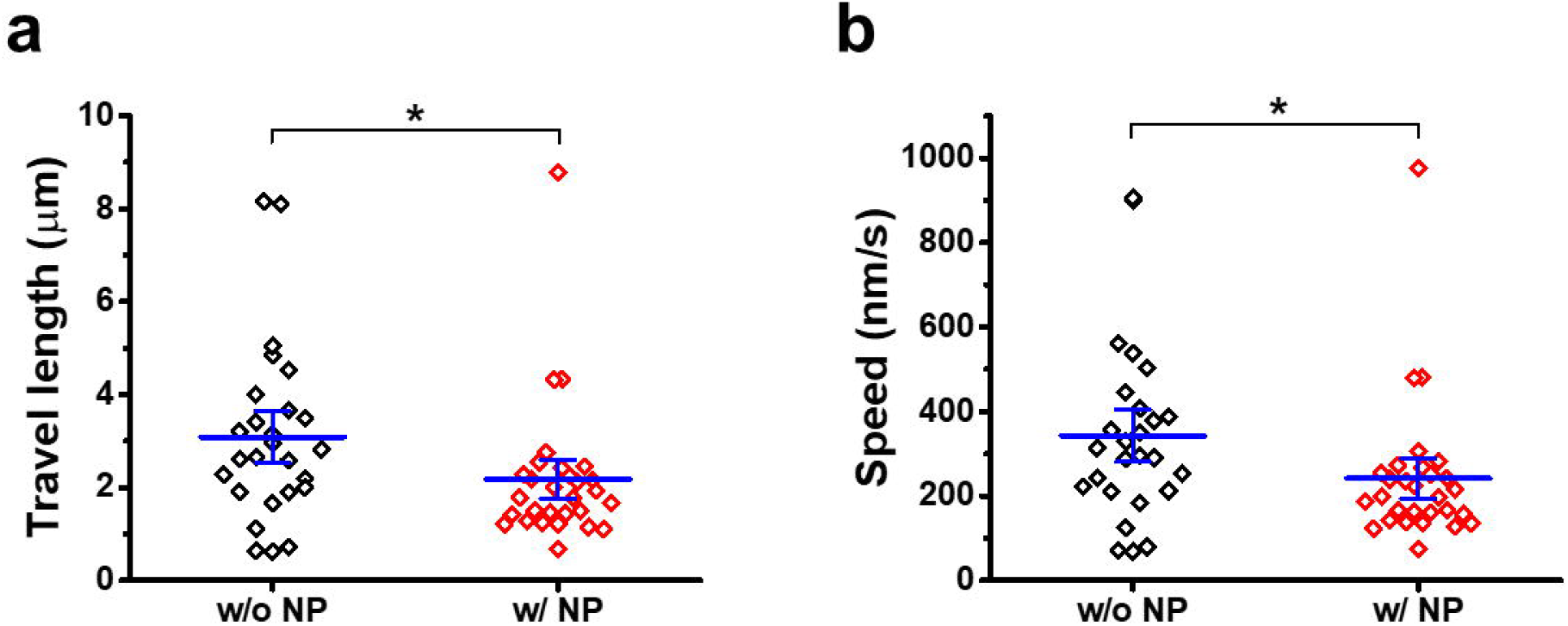
MPT-vVF reveals that exposure to nanoplastics impairs the transport of BDNF-containing vesicles in living neurons. a. Travel length of BDNF-containing moving vesicles in living neurons with and without 1 µg/mL yellow green 50 nm diameter polystyrene nanplastics. b. Speed of BDNF-containing moving vesicles in living neurons with and without 1 µg/mL nanplastics. \**p* < 0.05 (Mann– Whitney *U* test).

## Discussion

Accurate particle tracking is essential for understanding the spatiotemporal processes that underlie biological function(11, 16, 35, 36, 73, 74). Particle tracking has become an indispensable tool for quantitative investigations of biological dynamics(46, 74, 75). Quantitative analysis of motility, interaction, and intracellular transport based on particle tracking provide critical insights into the mechanisms governing cellular organization, signaling, and disease pathology(11, 12, 16, 20, 22, 26, 73, 76–79). However, accurate single particle tracking in living cells remains challenging. Intracellular environments are often densely populated with particles that exhibit heterogeneous motion behaviours, leading to frequent particle overlap, ambiguous particle associations, and trajectory fragmentation. The challenge is further compounded by the coexistence of stationary and motile particles, as signals from immobile structures can mask or distort the trajectories of nearby moving particles, thereby increasing the likelihood of tracking errors.

In this paper, we introduce MPT-vVF, a mathematically principled framework designed for robust tracking of moving particles in living cells. MPT-vVF exploits motion-induced spatiotemporal signatures to inherently suppress static background structures, thereby enhancing the contrast of moving particles without relying on extensive preprocessing. MPT-vVF employs two analytically derived filters to estimate instantaneous velocity with high accuracy. These velocity estimates serve as physically informed priors during trajectory linkage, reducing mislinkage in dense or crowded environments. MPT-vVF can accurately resolve nanometer-scale displacements of immobilized beads, as validated by precise tracking of fiducial fluorescent beads under controlled conditions. Critically, MPT-vVF successfully captures the dynamics of biologically relevant cargos, including single BDNF-containing vesicles in living neurons. Furthermore, MPT-vVF reveals that exposure to 50-nm nanoplastics impairs vesicular transport, reducing both travel length and speed of BDNF-containing vesicles in hippocampal neurons.

MPT-vVF addresses several limitations of existing methods for organelle tracking in living cells. Kymograph is widely used to characterize organelle transport in living cells(80) but requires substantial labour-extensive work including manual selection of particles and placement of lines along transport pathway. Moreover, the spatial resolution of kymograph is limited by the pixel size of the original image, which was 158 nm in our imaging system, making it difficult to resolve subpixel displacement. These constraints in kymograph may reduce throughput and increase user-dependent variability. We also evaluated the performance of TrackMate, which is widely used for tracking single particles in living cells(43), using the video shown in Fig. 4. Particle detection was found to be highly sensitive to the choice of detection threshold. At low thresholds, TrackMate detected hundreds of objects, the majority of which corresponded to background fluctuations rather than BDNF-mRFP-containing vesicles. Increasing the threshold reduced false-positive detections but retained numerous stationary high-intensity particles while failing to detect dimmer moving BDNF-mRFP-containing vesicles. Furthermore, the high density of BDNF-containing vesicles along neurites frequently led to misassignment of trajectories among neighboring particles. Although some of these issues can be mitigated through manual correction, such post-processing is labor-intensive and difficult to scale to large datasets. Similar issues may arise in the widely adopted optimization framework introduced by Jaqaman and colleagues(47), which links particle detections through a linear assignment problem (LAP) to construct trajectories. This framework may be susceptible to incorrect trajectory assignments in densely populated intracellular environments where particles exhibit frequent encounters, overlap events, and heterogeneous motion behaviors. In contrast, MPT-vVF exploits motion-derived spatiotemporal information and velocity-based constraints to improve both particle detection and trajectory association. By incorporating instantaneous velocity estimates into the tracking process, MPT-vVF substantially reduces tracking ambiguities in crowded cellular environments and enables more reliable reconstruction of organelle trajectories. These capabilities were particularly evident for BDNF-containing vesicles in neuronal processes, where high particle density pose significant challenges for conventional tracking methods. Furthermore, MPT-vVF is high parallelizable both across independent tracking tasks and within velocity estimation procedures using direction and magnitude, offering significant potential for acceleration on modern GPU architectures.

Recent advances in particle tracking have increasingly incorporated machine learning and artificial intelligence approaches. Deep learning frameworks such as Social LEAP Estimates Animal Poses (SLEAP)(81) enable flexible multi-object tracking through top-down or bottom-up pose estimation, while cross-modality image restoration models like cross-trained CARE (XTC)(82) enhance tracking input quality by computationally “super-resolving” low-SNR *in vivo* data. Nellie circumvents segmentation errors entirely by decoupling tracking from object delineation via motion-capture-inspired markers(39). Meanwhile, specialized hardware systems such as 3D-TrIm(83) use real-time active feedback to follow rapidly diffusing particles in three dimensions. Despite these innovations, many deep learning–based approaches remain “black-box” systems with limited interpretability, require extensive annotated training data, and often lack generalizability across organelle types or imaging modalities. In contrast, MPT-vVF is based on explicit physical principles governing particle motion, providing a transparent and interpretable framework for trajectory reconstruction. Collectively, our results demonstrate that MPT-vVF provides a robust and accurate framework for quantitative analysis of intracellular transport in living cells.

The applicability of MPT-vVF extends well beyond the tracking of vesicles in living cells. By combining motion-based detection with velocity-guided trajectory reconstruction, the framework can be readily adapted to diverse particle-tracking problems involving dense populations, heterogeneous dynamics, and frequent interactions. Its scalability across spatial and temporal scales makes it suitable for applications ranging from subcellular transport to collective dynamics in larger biological and soft-matter systems. Together, these features position MPT-vVF as a versatile computational platform for quantitative motion analysis in complex dynamic environments.

## Acknowledgments

We thank the members of Park lab and Zhang lab for helpful discussion and comments. This work was supported by the Research Grants Council of Hong Kong (Grants 16102322 and N_HKUST648/24 to H.P.) and by The National Natural Science Foundation of China (NSFC) (grant 12371425 to H.Z.)

## Author Contributions

X.L., Z.F., K.H.H., C.P.W., H.Z., and H.P. designed the method. K.H.H., J.Z., and H. P. performed the experiments. X.L., Z.F., C.P.W., C.P., Y.C., H.F.J.W., and H.P. analyzed data. X.L., Z.F., K.H.H., C.P.W., H.Z., and H.P. discussed the results. Y.Y., H.Z., and H.P. supervised students. X.L., Z.F., K.H.H., C.P.W., J.Z., C.P., H.Z., and H.P. wrote the manuscript.

## Declaration of Interest

The authors declare no competing interests

## Data Availability Statement

Research data will be shared upon reasonable request.

## Notes

### Competing Interest Statement

The authors have declared no competing interest.

